# Nitrone-Trolox conjugate as an inhibitor of lipid oxidation: Towards synergistic antioxidant effects

**DOI:** 10.1101/323386

**Authors:** L. Socrier, M. Rosselin, A. M. Gomez Giraldo, B. Chantemargue, F. Di Meo, P. Trouillas, G. Durand, S. Morandat

## Abstract

1

Free radical scavengers like α-phenyl-*N-tert*-butylnitrone (PBN) and 6-hydroxy-2,5,7,8-tetramethylchroman-2-carboxylic acid (Trolox) have been widely used as protective agents in various biomimetic and biological models. A series of three amphiphilic Trolox and PBN derivatives have been designed by adding to the parent molecules both a perfluorinated chain and a sugar group in order to render them amphiphilic. In this work, we have studied the interaction of these derivatives with lipid membranes and how it correlates to their antioxidant properties.

The three derivatives form monolayers at the air/water interface. We next investigated the ability of each derivative to interact with 1,2-dilinoleoyl-sn-glycero-3-phosphocholine (DLPC) as well as their efficiency to inhibit the AAPH-induced oxidation of DLPC liposomes. The location of these derivatives in the membrane is a key parameter to rationalize their antioxidant efficiency. The derivative bearing both the PBN and the Trolox antioxidant moieties on the same fluorinated carrier exhibited a synergistic antioxidant effect by delaying the oxidation process. Molecular dynamics (MD) simulations supported the understanding of the mechanism of action, highlighting various key physical-chemical descriptors.

**Graphical abstract:** 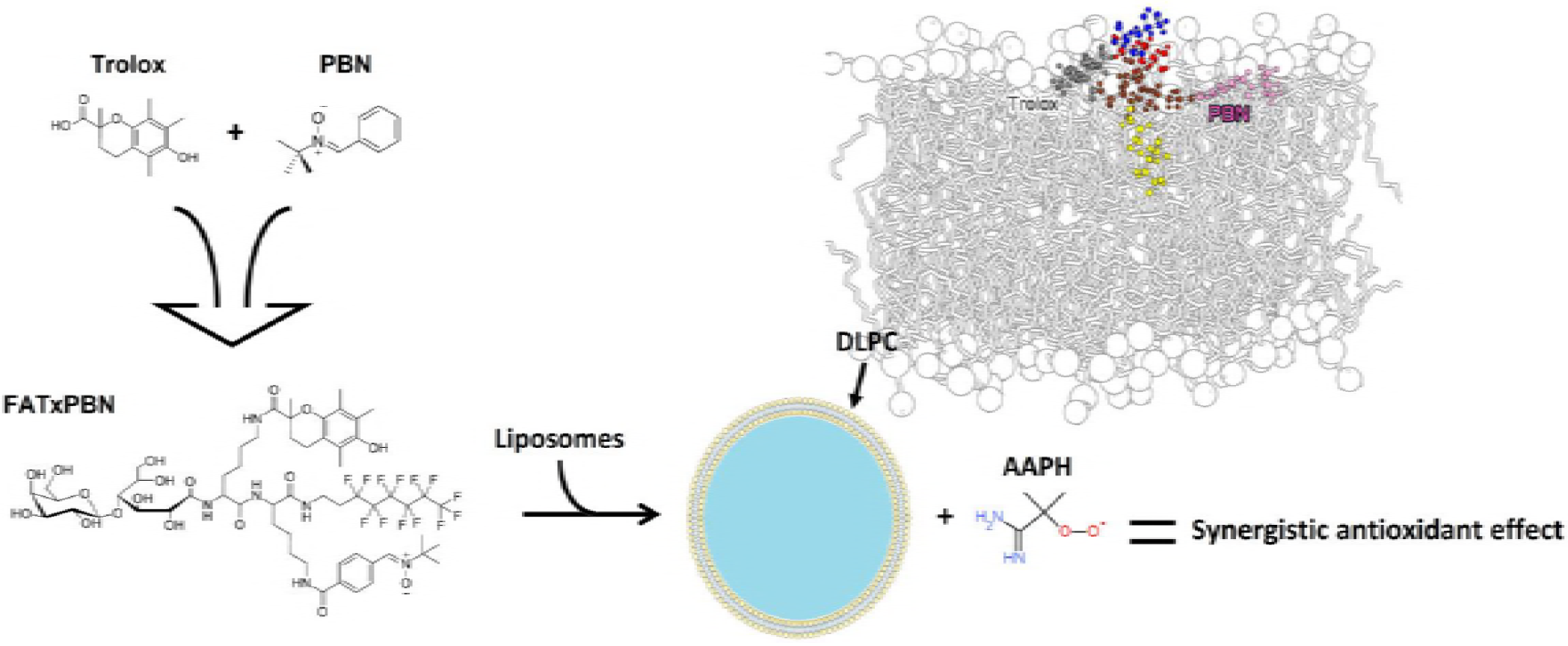

**Highlights:**

‒ Amphiphilic fluorinated antioxidants efficiently inhibit lipid oxidation
‒ The perfluorinated chain allows the insertion into membranes
‒ The nature of the antioxidant is a key parameter in the antioxidant efficiency
‒ The combination of Trolox and PBN results in a synergistic antioxidant effect

**Condensed running title:** Trolox derivatives limit lipid oxidation

## 1. Introduction

Reactive oxygen species (ROS) are essential in living organisms as they are implicated in several biological processes like signaling pathways (1), antimicrobial defense (2) or cell adhesion (3). However, deregulation of physiological processes as well as exposition to external factors including pollution, radiations and smoke (4) may favor ROS overproduction, which in turn can affect the oxidative stress status. Because of their ability to degrade biomolecules, ROS are involved in the early stages of several diseases like Alzheimer disease (5), diabetes (6), cell death (7) or cancers (8). Membrane lipids, especially polyunsaturated lipids (PUFAs), are particularly targeted by ROS to be degraded through a chemical process called lipid peroxidation (9).

To control the amount of ROS, living organisms use several antioxidant defense systems that are either endogenous (mainly enzymes but also small molecules as glutathione) or exogenous. Exogenous antioxidants (mainly polyphenols and vitamins) are basically natural and come from food. To improve their antioxidant properties, synthetic molecules have been developed based on these natural compounds. In this regard, Trolox^®^ (6-hydroxy-2,5,7,8-tetramethylchroman-2-carboxylic acid), a water soluble analog of α-tocopherol, has been used to prevent, *in vitro* or *ex vivo,* apoptosis (10) and ischemia (11). Trolox is also used as a reference in the evaluation of the antioxidant activity. Namely, the Trolox equivalent antioxidant capacity (TEAC) parameter measures the potency of a given antioxidant. TEAC also allows the comparison of antioxidant efficiency of food matrices and beverages, for which isolation of molecules can rarely been achieved (12, 13).

Because of their enhanced reactivity against free radicals, spin trapping agents are used as free radical scavengers (14). Among them, α-phenyl-*N-tert*-butylnitrone (PBN) is certainly the most popular. As most of spin-traps, PBN was first used in electron spin resonance (ESR) experiments, however, it has been used in biological models as an antioxidant (15, 16). Although PBN has the advantage of being a non-toxic chemically stable antioxidant with a half-life of six hours in animals (17), it presents a poor ability to remain inside cell membranes.

To create nitrones with higher hydrophobicity and affinity to membranes, several molecules deriving from nitrones have been designed. For instance, Ayuso *et al.* (18), as well as Choteau *et al.* (19, 20) reported the synthesis of PBN derivatives bearing a cholesterol moiety. In both cases, the authors demonstrated that compared to PBN, the derivatives presented an enhanced protective activity against stroke and retina light-induced damages, respectively. Along the same line, Trolox and PBN amphiphilic derivatives (21, 22) have recently been synthesized. They are bearing, on one hand, a perfluorinated chain to increase hydrophobicity without inducing a cytolytic effect (23) and on the other hand, a sugar group to maintain water solubility (Fig. 1). In the present work, we have studied the physical-chemical properties of these derivatives related to their capacity to interact with lipid membranes, which in turn has allowed unraveling their antioxidant properties by using biomimetic systems. Surface tension experiments showed the capacity of these derivatives to interact with lipids with respect to the parent (Trolox and PBN) derivatives. The antioxidant action at the membrane was evaluated by performing Langmuir monolayer and lipid peroxidation inhibition assays. The nature of the free-radical scavenger moieties and their location in the membranes were analyzed as key parameters to rationalize the antioxidant activity of these derivatives, which was supported by molecular dynamics (MD) simulations.

**Fig. 1.**
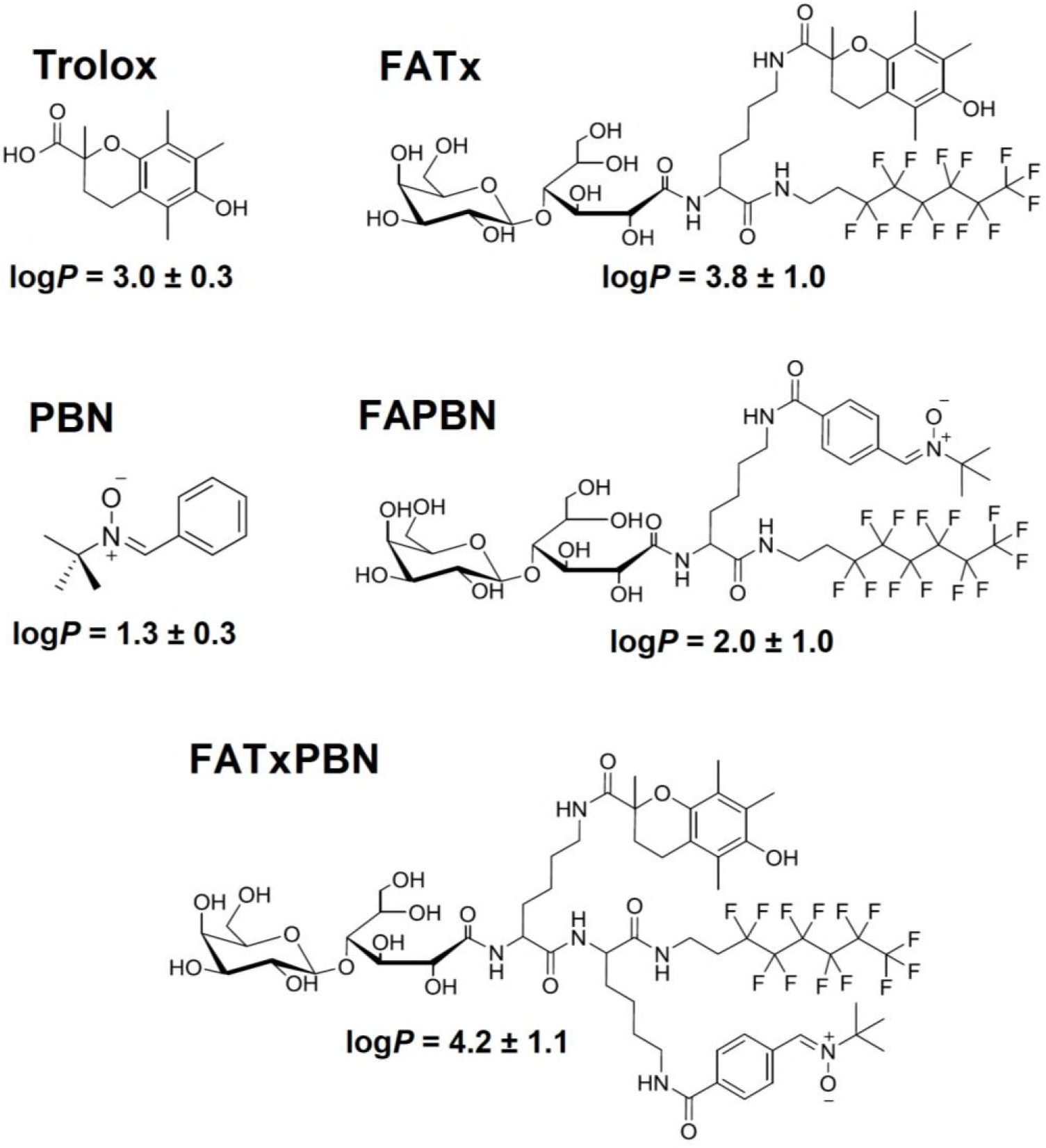
Chemical structures and log*P* values the studied molecules. Materials and methods.

## 2. Materials and methods

### 2.1. <u>Chemicals</u>

1,2-dilinoleoyl-*sn*-glycero-3-phosphocholine (DLPC), 1-palmitoyl-2-stearoyl-(5-DOXYL)-sn-glycero-3-phosphocholine (5-DOXYL PC) and 1-palmitoyl-2-steroyl-(10-DOXYL)-sn-glycero-3-phosphocholine (10-DOXYL PC) were purchased from Avanti Polar Lipids (Alabaster, AL, USA). 2,2′-azo-bis(2-amidinopropane)dihydrochloride (AAPH), 4-(2-hydroxyethyl)piperazine-1-ethanesulfonic acid (Hepes), (±)-6-hydroxy-2,5,7,8-tetramethylchromane-2-carboxylic acid (Trolox) and α-phenyl-*N-tert*-butylnitrone (PBN) were purchased from Sigma (St. Louis, MO, USA). FAPBN (21), FATx (21) and FATxPBN (22) were synthesized according to the procedure previously published.

The water used in all assays was purified using a Millipore filtering system (Bedford, MA, USA), yielding ultrapure water (18.2 MΩ×cm). Stock solutions of pure DLPC were prepared with chloroform or with hexane/ethanol 9:1 (v/v) for the preparation of liposomes or monolayers, respectively. The antioxidants were dissolved in chloroform or ethanol for the studies in liposomes or monolayers, respectively. log*P* values were calculated with the software ACD/ChemSketch version 14.01. www.acdlabs.com

### 2.2. <u>Surface tension experiments</u>

All the experiments were performed at constant temperature (21.0 ± 0.1°C). The film balance was built by Nima (Coventry, West Midlands, England) for adsorption experiments on a small Teflon dish. The dish (surface=19.6 cm^2^) was equipped with a Wilhelmy-type pressure measuring system. In these experiments, we used a subphase buffer of 62 mL continuously stirred with a magnetic stirrer spinning at a rate of 100 rpm. The buffer contained 10 mM of Hepes, 150 mM of NaCl and was adjusted at a pH of 7.4. To prepare Gibbs monolayers, the compounds - dissolved in ethanol - were injected in the subphase at two ranges of concentrations: from 1 to 40 μM for PBN and Trolox, from 0.0625 to 10 μM for the fluorinated derivatives. Surface tension was recorded during 5 min after the injection.

To study the adsorption of the compounds to DLPC monolayers, DLPC was dissolved in hexane/ethanol 9:1 (v/v) and spread at the air-water interface to reach the desired surface pressure. As soon as the initial surface pressure was stabilized (~ 15 min), the compounds were injected into the subphase at a final concentration of 50 nM. Their adsorption at the air-lipid buffer interface was then followed by measuring the variation of surface pressure. A control experiment was also performed by injecting ethanol in the subphase, which produced no significant perturbation of the surface pressure.

### 2.3. <u>Preparation of liposomes</u>

DLPC alone or DLPC/compound mixtures were dried under a stream of nitrogen and then kept under high vacuum for 2 hours to obtain a solvent-free film. The dry film was then dispersed in Hepes buffer to obtain multilamellar vesicles (MLVs). Small unilamellar vesicles (SUVs) were obtained by sonicating MLVs to clarity (3 cycles of 2 min 30 s) with a 500 W titanium probe sonicator at 33% of the maximal power from Fisher Bioblock Scientific (Illkirch-Graffenstaden, Grand-Est, France). To limit oxidation due to heat and light, sonication was performed in the dark and samples were kept in an ice bath. Then, the liposome suspension was filtered on 0.2 μm Acrodisc^®^ from Pall Life Sciences (Port Washington, NY, USA) to remove titanium particles. To prepare large unilamellar vesicles (LUVs), the MLVs suspension was extruded 19 times at room temperature through a 200 nm nuclepore polycarbonate membrane filter from Avestin Inc. (Ottawa, Ontario, Canada) with a syringe-type extruder from Avanti Polar lipids Inc. (Alabaster, AL, USA). Liposomes size distribution was determined by dynamic light scattering with a Zetasizer Nano-S from Malvern Instruments (Malvern, Worcestershire, UK) and was unimodal.

### 2.4. <u>Measurement of nitroxide quenching and evaluation of the depth inside the membrane</u>

DLPC Liposomes containing only 20 % (%mol) of fluorinated derivative or 20 % (%mol) of fluorinated derivative plus 15 % (%mol) of nitroxide labeled phospholipids (5-DOXYL PC or 10-DOXYL PC) were prepared with the procedure previously described (24). Fluorescence intensities were measured for each sample with the following excitation and emission wavelengths (λ_ex_:λ_em_); 345:687 nm for FAPBN, 300:595 nm for FATx and 255:505 nm for FATxPBN. The distance of the antioxidants from the center of the bilayer was calculated with the parallax equation (25):

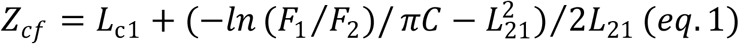

where Z_cf_ represents the distance of the derivative from the center of the bilayer, L_c1_ the distance of the shallow quencher from the center of the membrane, F_1_ the fluorescence intensity (F_5_/F_0_) in the presence of the shallow quencher (5-DOXYL PC), F_2_ the fluorescence intensity (F_10_/F_0_) in the presence of the deeper quencher (10-DOXYL PC), C the concentration of quencher in molecules/Å^2^. To be more precise, C corresponds to the mole fraction of nitroxide-labeled phospholipids divided by the area per phospholipids (70 Å), *i.e.*, 0.15/70 Å. At last, L_21_ corresponds to the distance between the shallow and deep quenchers. Fluorescence intensities of DLPC alone were verified and the values obtained were not significant.

### 2.5. <u>Measurement of the effect of the compounds on the permeability of the membrane</u>

The ability of the fluorinated derivatives to permeabilize liposomes was measured with the calcein release assay. DLPC LUVs were prepared as described above with Hepes buffer containing 35 mM calcein. Free calcein was separated from encapsulated calcein by gel filtration on a sepharose 4B column equilibrated with Hepes buffer. Lipids were quantified with the method developed by Stewart (26). The calcein loaded liposomes were diluted to a final concentration of 10 μM lipids in Hepes buffer. Fluorinated derivatives were added after 5 min at a ratio of 5 % (mol/mol) compared to DLPC. Fluorescence measurements were performed at 21 °C as soon as the liposomes were prepared, immediately after the addition of fluorinated derivatives and after 30, 60 min and 120 min of incubation with Varian Cary Eclipse fluorescence spectrophotometer (Santa Clara, CA, USA). We used excitation and emission wavelengths of 490 and 520 nm respectively. Triton X-100 was added after two hours in order to release the total amount of encapsulated calcein. The intrinsic fluorescence of fluorinated antioxidants at 520 nm was verified and the values we found were not significant. Release percentages were calculated with the following equation:

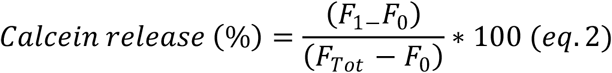

where F_1_ is the fluorescence of the liposomes measured with fluorinated derivatives, F_0_ the fluorescence of the liposomes measured in absence of fluorinated derivatives and F_Tot_ the fluorescence measured after addition of Triton X100 that releases the encapsulated calcein.

### 2.6. <u>Antioxidant efficiency of fluorinated derivatives</u>

The antioxidant efficiency of the non-modified derivatives as well as the derivatives was assessed by measuring the amount of conjugated dienes, which are primary peroxidation products for PUFAs (9, 27). Liposomes of DLPC alone or containing fluorinated derivatives, Trolox or PBN at different concentrations were prepared in Hepes buffer. Lipid peroxidation was initiated by adding AAPH at a final concentration of 2 mM. The final concentration of DLPC for each assay was kept constant at 0.1 mg/mL. The samples were immediately incubated at 37°C. The formation of conjugated dienes was followed by measuring the absorbance at 234 nm with a Specord S300 UV-VIS spectrophotometer (Jena, Thuringia, Germany). The values were normalized to zero by subtracting the initial absorbance (0.8 ± 0.2). For the control assay, DLPC liposomes were tested alone. The results are expressed as percentage of peroxidation, calculated as follows:

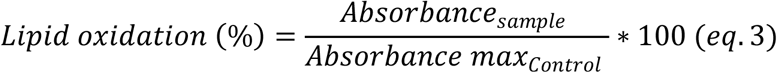

The curves were fitted by applying a dose-response model with variable slope with the software GraphPadPrism version 6.00 for Windows. (La Jolla, CA, USA). www.graphpad.com.

To further characterize the differences in the antioxidant efficiency of the derivatives, we calculated the lag time of oxidation as follows:

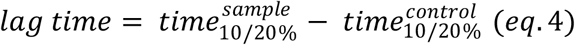

where *time*_*10/20%*_ corresponds to the time required to reach 10 or 20 % of oxidation for the DLPC liposomes containing 6,4 μM of antioxidant (5 %, mol/mol compared to DLPC) (sample) and for the pure DLPC vesicles (control).

### 2.7. <u>Theoretical methodology</u>

Two DLPC membranes made of 128 and 72 lipids each, were created using the membrane bilayer builder from the CHARMM-GUI server (28). The membranes were solvated with a hydration number of 35 water molecules per lipid. Na^+^ and Cl^−^ ions were added at a 0.154 M concentration, ensuring the neutrality of the system. The DLPC lipids were described using the lipid 11 force field (29). The force field parameters for the different derivatives (Trolox, anionic Trolox, PBN, FATx, FAPBN, FATxPBN) were derived from the Generalized Amber Force Field version 2 (GAFF2) (30–32) for all moieties but the sugars, which were described by the GLYCAM force field (33), using the antechamber package (34). The FATx, FAPBN, FATxPBN were built using a building block approach (*i.e.*, considering the β-D-galactopyranosyl, 2,6-diaminohexanoyl, 1-amino-dodecafluorooctanyl, Trolox and PBN residues). Atomic charges were derived from RESP (Restrained fit of Electro Static Potential) based on the calculations achieved within the Density Functional Theory (DFT) formalism with the IEFPCM-B3LYP/cc-pVDZ method in diethylether (35), accounting for the chemical environment in the building block approach. The DFT calculations and the atomic charge fitting were performed with the Gaussian 09, RevA (36) and R.E.D. III (37) softwares, respectively. The three-point TIP3P water model (38) was used to describe water molecules.

MD simulations were carried out using CPU-Particle-Mesh Ewald (PME) (39) MD codes available in Amber16 (40, 41), and according to the following procedure: minimization of water molecules prior to minimization of the entire system to prevent from steric clash; slow thermalization of the water molecules up to 100 K in the (N,V,T) ensemble for 200 ps; thermalization of the whole system to the final temperature (310 K) for 500 ps in the (N,P,T) ensemble; equilibration of the system for 5 ns (N,P,T) MD simulations. Productions of 600 ns (N,P,T) MD simulations were then achieved. PME MD simulations were carried out using the SHAKE algorithm (42) on H-bonds and a 10 Å non covalent interaction cut-off for both Coulomb and van der Waals interactions. The temperature was maintained using the Langevin dynamics (43) with a collision frequency of 1 ps^−1^. Semi-isotropic pressure scaling with constant surface tension in the xy-plan perpendicular to the membrane normal (z-axis) was used with Berendsen barostat (44), in which the pressure relaxation time was set to 1 ps.

The different derivatives were inserted in the bulk water of the equilibrated membrane systems, preventing from steric clash with water molecules. MD simulations of 600 ns were then carried out with all derivatives; the total MD simulation time for the six derivatives was 3.6 μs. The analyses were performed along the last 300 ns, which allowed a robust sampling of structural properties, *i.e.*, after the equilibrium was reached. They were carried out using the cpptraj software (45). The depth of insertion of the different derivatives was measured from the center of mass (COM) of the lipid bilayer (*i.e.*, z-component of the vector originated at the COM of the lipid bilayer and pointing towards the COM of the derivative). The orientation of the derivatives in the lipid bilayer was assessed by the α-angle between the z-axis and the normal vector(s) defined for each derivative (see supplemental data).

## 3. Results and discussion

### 3.1. <u>Interfacial behavior of the antioxidants</u>

#### 3.1.1. Adsorption of the fluorinated derivatives at the air-buffer interface

To characterize the amphiphilicity of the Trolox and PBN derivatives, we measured their ability to adsorb at the air-buffer interface. Increasing concentrations were injected in the subphase and the surface tension was recorded continuously to assess the formation of a Gibbs monolayer at the interface. Fig. 2 represents the surface activity of the fluorinated derivatives. The injection of 1, 5, 10, 20 and 30 μM of Trolox and PBN exhibited only a poor effect on the surface pressure as the variations of surface pressure (Δπ) recorded were ranging between 0 and 3 mN/m (data not shown). Conversely, the fluorinated derivatives presented a high surface activity since the injection of 0.1 μM was sufficient to observe a significant surface activity (Fig. 2A–C). A maximal surface activity of ~ 40 mN/m was reached at only 1 μM for FATxPBN and FATx and at 3 μM for FAPBN, indicating that both FATxPBN and FATx are more hydrophobic than FAPBN. Higher concentrations (*i.e.* 5 and 10 μM) were also tested and the Δπ values remained close to 40 mN/m.

The log*P* values of the derivatives were calculated and are summarized in Fig. 1. The three fluorinated derivatives exhibit log*P* values significantly higher than their parent derivatives. This confirmed the strong contribution of the perfluorinated chain to the overall hydrophobicity of amphiphilic molecules. Moreover, the higher hydrophobicity of Trolox over PBN is clearly observed, with log*P* values of 3 and 1.3, respectively. This is also observed when comparing FATx and FAPBN with log*P* values of 3.8 and 2, respectively. The same tendency was observed with the experimentally-determined log*k′*_w_ values previously reported (22). FATxPBN and FATx exhibit log*k′*_w_ values ~ 5 times higher than Trolox when that of FAPBN is 3 times higher than PBN. The higher hydrophobic character of the fluorinated derivatives compared to the parent derivatives agrees with the pronounced surface activity, reflecting the formation of stable monolayers at the air-water interface.

**Fig. 2.**
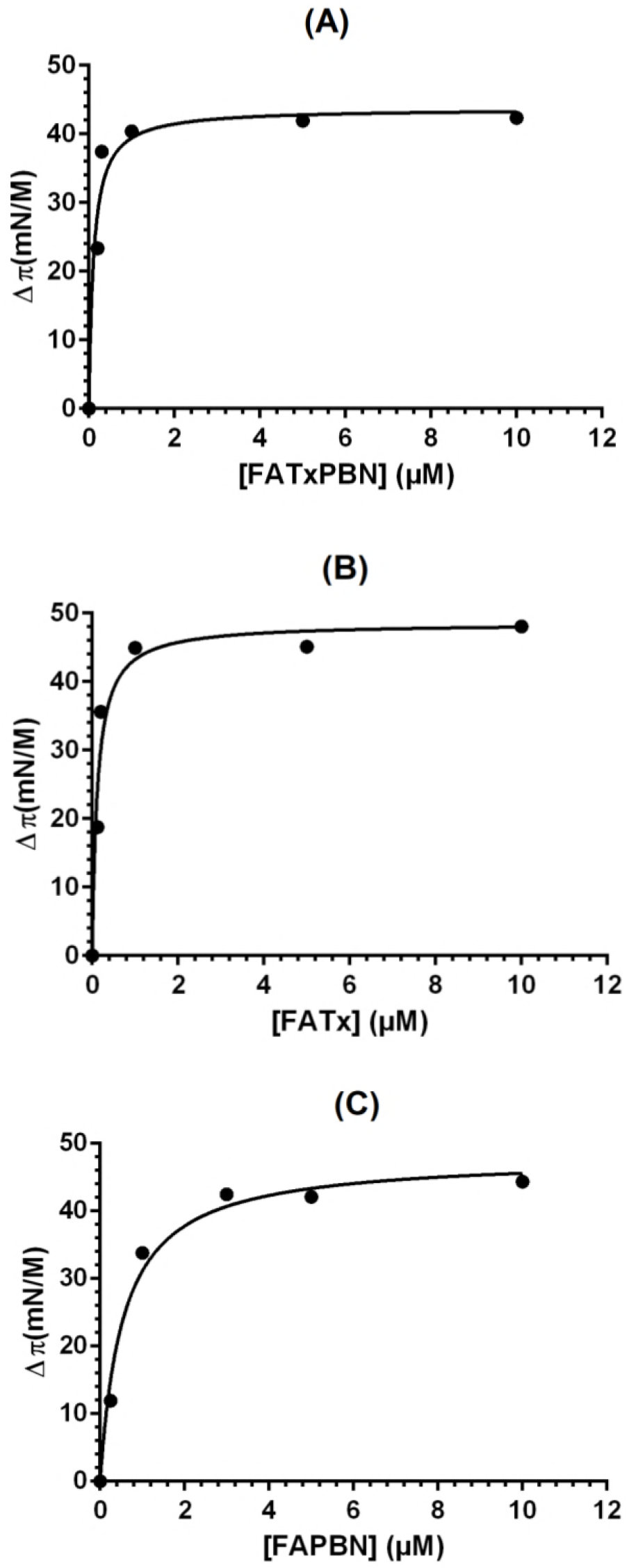
Surface activity (Gibbs monolayers) of the fluorinated derivatives at the air/water interface. FATxPBN (A), FATx (B) and FAPBN (C). Data were fitted with a Langmuir equation with GraphPadPrism software.

#### 3.1.2. Interactions with an unsaturated phospholipid

To evaluate the ability of each derivative to interact with DLPC, we performed assays of adsorption at lipid monolayers. DLPC monolayers were prepared at different initial pressures and the derivatives were injected in the subphase. The variation of pressure was recorded to evaluate the ability of the derivatives to modify the surface activity of the monolayer. To avoid that the own surface activity of the fluorinated derivatives affects the measurements, they were injected at low concentration (0.05 μM) since at concentrations lower than 0.1 μM they all presented a negligible surface activity (see Fig. 2). Fig. 3 represents the variation of pressure (Δπ) *versus* the initial surface pressure (π_0_). The injection of the parent compounds had little effect on the surface pressure, as Δπ remains below 3 mN/m independently of the initial pressure (data not shown). Conversely, the injection of the fluorinated derivatives (Fig. 3A–C) caused an immediate increase of the surface pressure and Δπ decreased when the initial pressure was increased. Linear regression provided the maximal insertion pressure (MIP), highlighting the highest pressure at which fluorinated derivatives can penetrate monolayers (46). As reported on Fig. 3, FATxPBN, FATx and FAPBN exhibit MIPs of 51.5, 48.6 and 39.5mN/m, respectively. Because all the MIPs are higher than 30-35mN/m, which have been reported to be the average lateral pressure in biological membranes (47). This suggests that fluorinated derivatives can penetrate bilayers under physiological pressure. This result shows that the addition of the fluorinated chain has increased the affinity of the parent compounds for DLPC.

Calvez and co-workers (46) reported that in addition to MIP, another parameter can be used to evaluate the affinity of the fluorinated derivatives to partition in DLPC. Indeed, the extrapolation from an initial surface pressure of 30 mN/m allowed to determine Δπ_30mN/m_, which reflects the quantity of fluorinated derivatives associated to the monolayer. As reported in Fig. 3, FATxPBN and FATx exhibit Δπ_30_ values ~ 4 times higher than FAPBN (9.9 and 9.0 versus 2.3 mN/m). This result indicates that the amount of FATxPBN and FATx inserted in the membrane is much more important than the amount of FAPBN.

**Fig. 3.**
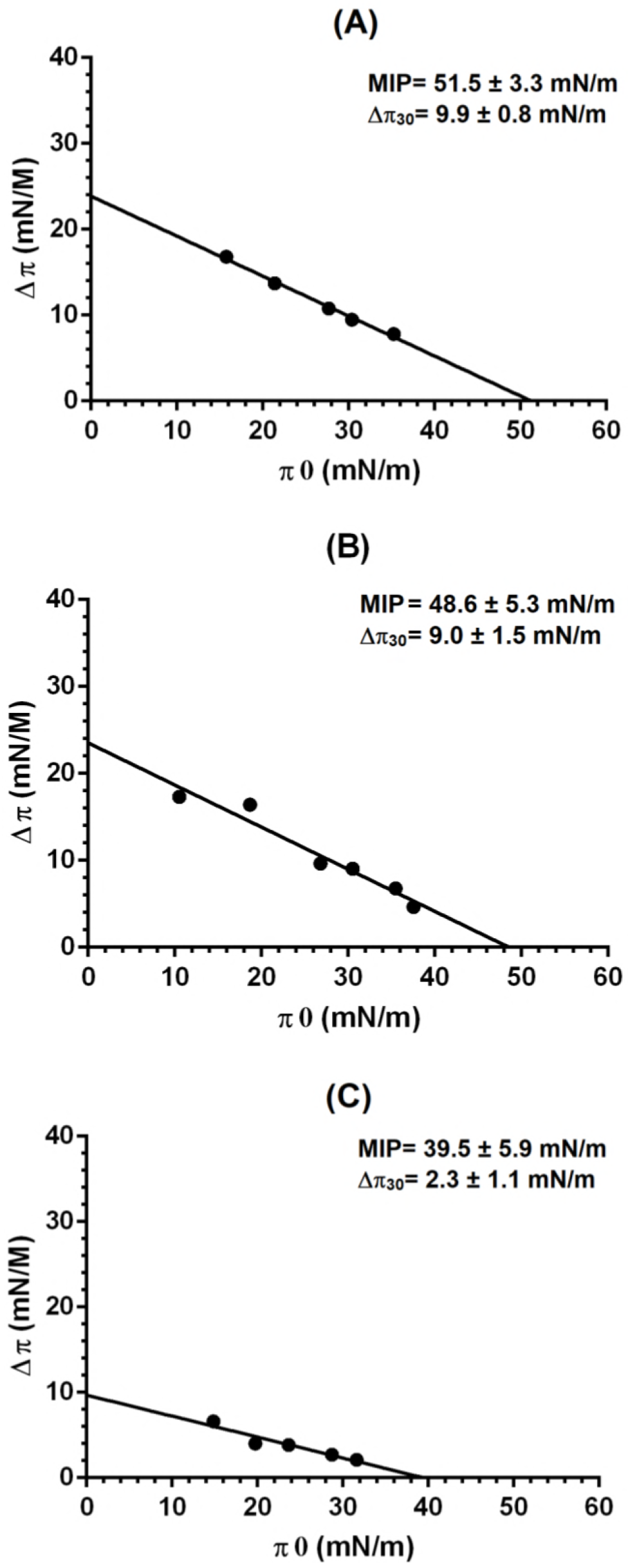
Interactions of the fluorinated derivatives with DLPC monolayers. FATxPBN (A), FATx (B), FAPBN (C). Upper right: Maximal insertion pressure (MIP) and Δπ_30 mN/m_ values of the fluorinated derivatives.

### 3.2. <u>Hydrophobicity affects the insertion of the derivatives in membranes</u>

The three fluorinated derivatives were shown capable to penetrate DLPC monolayers. However, differences in affinity were observed and were correlated to the overall hydrophobicity. The parallax method (25) allowed estimation of the position of the derivatives in the DLPC membrane.

DLPC liposomes containing the fluorinated derivatives and a nitroxide-labeled lipid were prepared. Fluorescence intensities were measured, from which the distance of each derivative from the center of the membrane (Z_cf_) was estimated. Distances of the nitroxide group from the center of the membrane were 12,2 Å and 6,1 Å for 5-DOXYL PC and 10-DOXYL PC, respectively (25). As seen from Z_cf_ values in Table 1, both FATxPBN and FATx appear embedded deeper inside the membrane than FAPBN. The log*P* values confirmed that the higher the hydrophobicity, the deeper the molecules (Supp. 1). These results agree with the Δπ_30mN/m_ values, suggesting that FAPBN has less affinity to the membrane than the other derivatives.

**Table 1.** Distance Z_rf_ of the fluorinated derivatives from the center of the membrane.

|  | **FATxPBN** | **FATx** | **FAPBN** |
| --- | --- | --- | --- |
| **$Z_{cf}$ (Å)** | 8.7 ± 0.1 | 9.2 ± 0.2 | 12.3 ± 1.0 |

### 3.3. <u>The fluorinated derivatives do not modify the membrane’s structure</u>

We performed calcein release experiments to evaluate the impact of the fluorinated derivatives on the membrane integrity. DLPC liposomes containing calcein were prepared and the fluorinated derivatives were added to the samples while fluorescence was measured during 2 hours to evaluate the calcein leakage. In all cases, the addition of the derivatives caused a weak leakage from the liposomes, with however a percentage of leakage below 5 % during the entire experiment (Supp. 2). The addition of Triton X100 after two hours caused a significant increase of the fluorescence intensity due to the release of the total amount of encapsulated calcein. Because the injection of the derivatives did not significantly modify calcein leakage, we concluded that their insertion has no significant effect on the structure of the membrane. Frotscher and co-workers (48) observed a solubilization of lipid bilayers after incubation of a fluorinated surfactant with a similar polar head group. Our assays did not reveal any significant modification of the permeability of the membrane, indicating that the fluorinated derivatives tested here did not solubilize the membrane. This is very likely due to the very low concentration (physiologically more relevant) used in the present study (5 μM) compared to the 2 mM used by Frotscher and co-workers.

### 3.4. <u>Fluorinated derivatives are efficient lipid peroxidation inhibitors</u>

Previous experiments demonstrated that Trolox and PBN have less affinity to the membrane than their fluorinated derivatives, which can penetrate DLPC monolayers. In order to evaluate the antioxidant efficiency of the compounds, DLPC liposomes containing the parent compounds or the fluorinated derivatives were prepared and incubated with AAPH. Oxidation percentages were calculated with eq.3 as previously described and reported over time. In absence of antioxidant, the percentages of oxidation increase until reaching a plateau at 60 min (Fig. 4A). This result stands from the focus given to the conjugated dienes which are the primary peroxidation products (9, 27), and which in turn drive the beginning of the kinetics. Consequently, after 80 min, the percentage of lipid oxidation decreases, while oxidation continues beyond the conjugated dienes stage. Namely, conjugated dienes are progressively replaced by secondary peroxidation products that are not detected at 234 nm. With increasing concentrations of FATxPBN, we noticed a gradual diminution of the percentages, showing its dose-dependent antioxidant efficiency (Fig. 4A). The same tendency was observed with Trolox, FATx and FAPBN (data not shown).

To better compare the antioxidant efficiencies, we reported the percentage of oxidation at 80 min (*i.e.*, the time required to reach the plateau in the absence of antioxidant) versus the antioxidant concentration (Fig. 4B–F). For FATxPBN and FATx, concentrations ranging from 1.6 to 6.4 μM appear sufficient to reach the maximal antioxidant efficiency. FAPBN is significantly less efficient as saturation was reached with concentrations ranging from 25 to 38 μM. No significant antioxidant efficiency was observed for PBN, even at high concentrations (*i.e.* 30-40 μM). Trolox, FATxPBN, FATx and FAPBN present IC_50_ (concentration required to half-reduce the oxidation in absence of antioxidant) values of 4.0, 0.6, 0.9, and 29.5 μM, respectively, no IC_50_ was determined for PBN.

The better lipid peroxidation inhibitor capacity of Trolox compare to PBN agrees with previous work, where we showed that Trolox was more efficient than PBN to scavenge ABTS^+•^ radicals (22). All three fluorinated derivatives exhibit lower IC_50_ than their parent derivatives. Indeed, FATxPBN and FATx exhibit IC_50_ ~ 5 times lower than Trolox (0.6 and 0.9 μM versus 4.0 μM) and FAPBN takes an IC_50_ of 29.5 μM while PBN has no IC_50_.

**Fig. 4.**
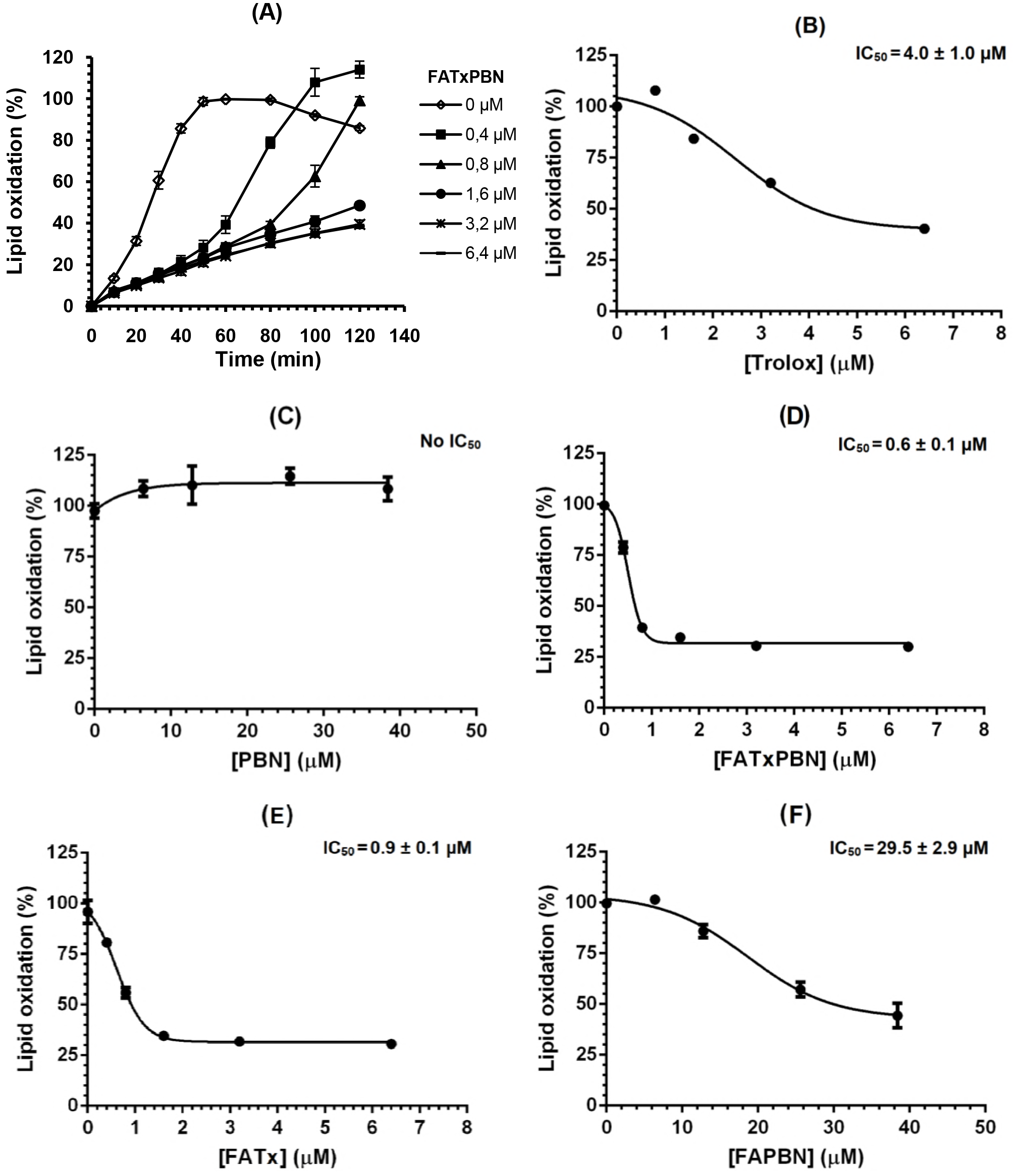
Effect of the parent compounds and the fluorinated derivatives on the AAPH-induced oxidation of DLPC liposomes. Kinetics of oxidation in presence of FATxPBN (A). IC_50_ (μM) of Trolox (B), PBN (C), FATxPBN (D), FATx (E) and FAPBN (F). Results are presented as the mean ± standard deviation of at least three experiments.

To further discriminate antioxidant efficiencies of the most two efficient derivatives (FATxPBN and FATx), we determined the lag times, which represent the time required to reach either 10 or 20% lipid peroxidation for DLPC liposomes containing the same amount of the different antioxidants, *i.e.* 6,4 μM (Fig. 5). Except PBN and FAPBN at 20% oxidation (1.5 ± 2.0 and 6.0 ± 1.0 min, respectively), the lag times of the fluorinated derivatives do not significantly differ compared to that of the parent derivatives. In the case of FATxPBN, the lag time was significantly higher at 10 and 20% oxidation than for all other derivatives, which suggests that this molecule has a different kinetic inhibition process and it efficiently delays the oxidation process. Along the same line with the rest of this work, these data confirmed a significantly improved antioxidant efficiency of the fluorinated derivatives with respect to the parent derivatives.

**Fig. 5.**
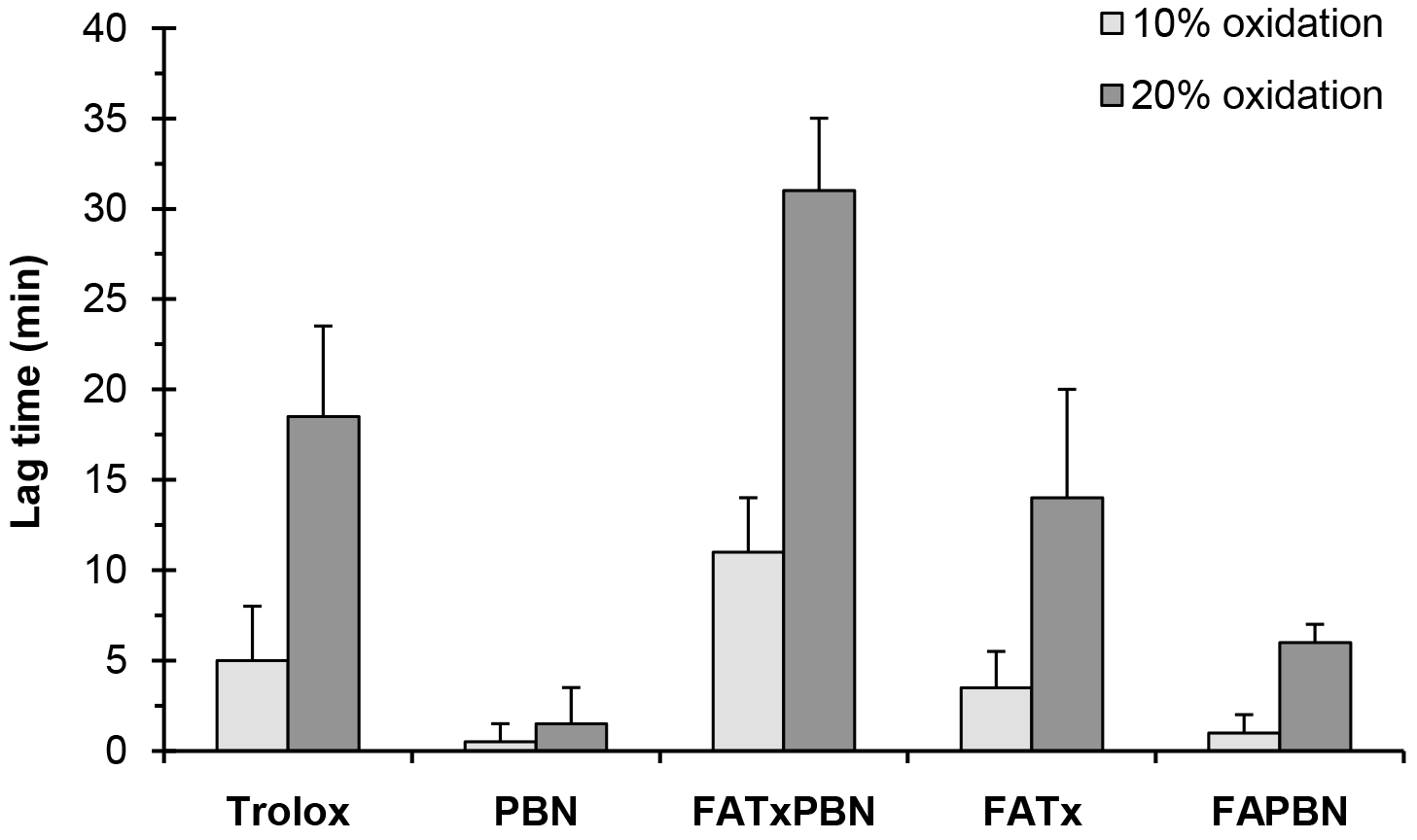
Lag time necessary to reach 10 and 20 % of oxidation. Results are presented as the mean ± standard deviation of at least three experiments.

### 3.5. <u>MD simulations of derivative positioning in DLPC membrane</u>

All derivatives inserted into the DLPC lipid bilayers when starting from the bulk water. The preferred position of the center of mass (COM) of Trolox and PBN were just below the polar head group region in close contact with the phosphates and to a less extent with the choline moieties (Table 2 and Supp. 3 for the radial distribution function). The fluorinated derivatives also partition just below the polar head group region. The different moieties drive the preferred location of these amphiphilic derivatives. Namely, from one side, both *β*-D-Galactose and gluconic acid moieties strongly anchor the derivatives to the choline and the phosphate groups by electrostatic and H-bonding interactions (Fig. 6). From the other side, the fluorinated chain lines up along the DLPC lipid chains, anchoring the derivatives into the membrane (Fig. 6). In the fluorinated derivatives, the Trolox and PBN moieties fluctuate below the polar head group region with a preferred orientation parallel to the surface (Supp. 4) in close contact with the phosphates, as for the parent Trolox and PBN derivatives. At this preferred location, *i.e.*, in contact with the lipid/water interface, the derivatives are likely to inhibit both the initiation stage or the propagation stage for lipid chains which can adopt a transient snorkel-like shape (49, 50) of lipid peroxidation. Interestingly, in the case of FAPBN, a minor arrangement was observed inside the membrane, which drives the PBN moiety deeper in the membrane (Table 2), in close contact with the fluorinated chain (Fig. 6D). In this arrangement, inhibition of the propagation stage is likely to occur. A similar conformational rearrangement is also expected in FATxPBN, *i.e.*, lining up the PBN moiety along the fluorinated chains. Conversely, similar alignment of the Trolox moiety and the fluorinated chain (in FATx and FATxPBN) has appeared unlikely, even though we have conducted a MD simulation in which these two moieties were in closer contact. It is worth noting that no intramolecular interactions between the Trolox and PBN moieties were observed in FATxPBN throughout the whole MD simulation.

**Fig. 6.**
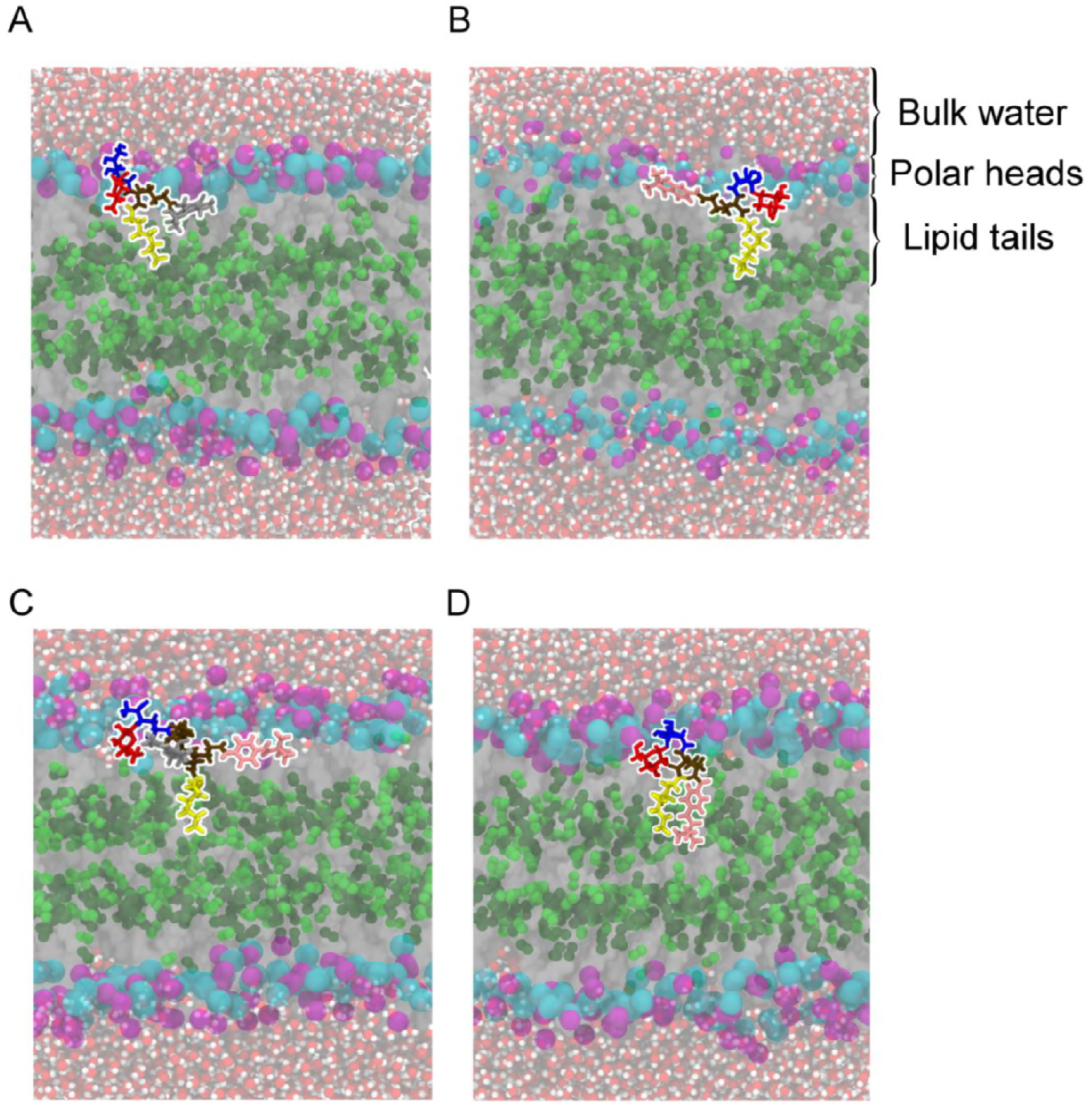
Representative snapshots of the preferred location of FATx (A), FAPBN (B) and FATxPBN (C). Fig. 6D depicts the minor conformation of FAPBN, in which the PBN moiety is lined up along the fluorinated chain. The 1LB (β-D-Galactose), GA4 (gluconic acid), DAH (2,6-diaminohexanoic acid) and DFA (3,3,4,4,5,5,6,6,7,7,8,8,8-dodecafluorooctan-1-amine) moieties are depicted in red, blue, brown and yellow, respectively. Trolox and PBN are depicted in grey and pink, respectively.

**Table 2.** Position (in Å) of the key moieties of the DLPC (*i.e.*, choline, phosphates, unsaturations) and the different derivatives Trolox, Anionic Trolox, PBN, FATx, FAPBN and FATxPBN (*i.e.*, Trolox moiety, Trolox OH-phenolic, PBN, PBN amine moiety, 1LB = β-D-Galactose, GA4 = gluconic acid; DAH = 2,6-diaminohexanoic acid, DFA = 3,3,4,4,5,5,6,6,7,7,8,8,8- dodecafluorooctan-1-amine), as well as center-of-mass (COM) of the three fluorinated derivatives FATx, FAPBN and FATxPBN. The positions were averaged over the second half of the MD simulations.

|  | Trolox | Anionic Trolox | PBN | FATx | FAPBN | FATxPBN |
| --- | --- | --- | --- | --- | --- | --- |
| **Cholines** | 20.4 ± 0.4 | 20.3 ± 0.4 | 20.5 ± 0.4 | 20.5 ± 0.3 | 20.5 ± 0.3 | 20.5 ± 0.3 |
| **Phosphates** | 19.1 ± 0.4 | 19.0 ± 0.4 | 19.2 ± 0.3 | 19.2 ± 0.3 | 19.2 ± 0.2 | 19.2 ± 0.3 |
| **1<sup>st</sup> lipid unsaturation** | 7.2 ± 0.3 | 7.2 ± 0.3 | 7.2 ± 0.3 | 7.2 ± 0.2 | 7.3 ± 0.2 | 7.2 ± 0.2 |
| **2<sup>nd</sup> lipid unsaturation** | 5.6 ± 0.2 | 5.6 ± 0.2 | 5.7 ± 0.2 | 5.7 ± 0.2 | 5.7 ± 0.1 | 5.7 ± 0.2 |
| **Trolox** | 14.0 ± 1.7 | 15.3 ± 1.8 | | 14.0 ± 2.3 | | 13.9 ± 1.7 |
| **Trolox phenolic OH group** | 15.1 ± 2.2 | 14.2 ± 2.2 | | 14.7 ± 2.6 | | 14.0 ± 2.3 |
| **PBN** | | | 13.6 ± 2.7 | | 15.7 ± 2.5 | 13.8 ± 1.5 |
| **PBN amine** | | | 13.8 ± 2.8 | | 15.2 ± 2.9 | 13.9 ± 1.8 |
| **1LB** | | | | 16.7 ± 1.7 | 16.3 ± 2.0 | 16.8 ± 1.9 |
| **GA4** | | | | 20.4 ± 1.5 | 20.0 ± 1.9 | 19.6 ± 1.7 |
| **DAH** | | | | 16.3 ± 1.9 | 16.2 ± 1.8 | 15.2 ± 1.2 |
| **DFA** | | | | 10.0 ± 1.8 | 10.4 ± 1.9 | 8.2 ± 1.6 |
| **FATx COM** | | | | 14.3 ± 1.7 | | |
| **FAPBN COM** | | | | | 14.7 ± 1.7 | |
| **FATxPBN COM** | | | | | | 13.7 ± 1.1 |

### 3.6. <u>Mechanism of lipid peroxidation inhibition of FATx, FAPBN and FATxPBN</u>

The antioxidant assays conducted in this work highlight that the three fluorinated derivatives are much more efficient than their parent derivatives. However, significant discrepancies were observed between the derivatives, as both FATxPBN and FATx exhibit IC_50_ that are 49 and 33 times lower (higher activity) than FAPBN, respectively (Fig. 4). The differences observed between FAPBN and the other two fluorinated derivatives can be rationalized by a lower affinity to lipid (slightly lower hydrophobicity, see log*P* on Fig. 1) of the former derivative and by the absence of the Trolox moiety in this derivative. The scavenging mechanisms of both Trolox and PBN are different (most probably hydrogen atom abstraction and adduct formation - spin trap -, respectively), which makes the former derivative a much better free radical scavenger. Concomitantly, the amphiphilic feature of the derivatives finely tunes the capacity for lipid peroxidation inhibition, by ensuring a positioning in the membrane in close contact with the oxidation process (*i.e.*, as close as possible from unsaturation). It is well-known that location of the antioxidants in close contact to the unsaturations of the phospholipid acyl chains favors a better inhibition of lipid peroxidation (51). For instance, Azouzi and co-workers recently reported that serotonin has the ability to bind and protect DLPC enriched membranes from oxidation (52), due to frequent interactions between serotonin and phospholipid acyl chains, which might inhibit the propagation stage of lipid peroxidation.

Interestingly, PBN exhibits no lipid peroxidation inhibition whereas an IC_50_ was measured with FAPBN. Although both PBN itself and the PBN moiety of FAPBN lie at almost the same preferred position, one can expect an anchoring effect attributed to the fluorinated chain which better constrain (anchor) the PBN moiety positioning in the membrane, which can more efficiently act as a spin trap. In its minor FAPBN conformation (Fig. 6D), the PBN moiety may even spend some time deeper in the bilayer to inhibit lipid peroxidation more efficiently than PBN alone, however not as efficient as both FATxPBN and FATx due to a much less efficient free radical scavenging capacity.

FATx is 33 times more active than FAPBN, which is mainly attributed to the nature of the antioxidant moiety and not to positioning. Indeed, Trolox is a much more efficient free radical scavenger than PBN, while both moieties are positioned in the same membrane region, in close contact with the polar head group region. With an IC_50_ of 0.6 μM, FATxPBN has shown the highest antioxidant activity among all three fluorinated derivatives. Because its IC_50_ is smaller than the average of FATx and FAPBN, we suggest that the presence of Trolox and PBN on the same carrier creates a synergistic antioxidant effect. To evaluate the presence of such an effect, we focused on the behavior of FATx and FAPBN with respect to that of FATxPBN. Both FATx and FAPBN were considered here as two separated antioxidant agents engaged in a similar skeleton, *i.e*., an antioxidant moiety linked to a perfluorinated chain and a sugar moiety. FATxPBN combines these two antioxidant agents, and this combination can result either simply in additive, or in supra-additive (or synergistic), or in infra-additive (inhibition of one agent by the other) effects. To do so, the percentage of lipid oxidation of FATx and FAPBN were multiplied and compared to FATxPBN (53, 54). As one can see in Table 3, at very low concentrations, (*i.e.*, 0.4 to 0.8 μM), FATxPBN presented a synergistic effect since its percentages of oxidation are lower than the multiplicative combination of FATx and FAPBN. This effect diminishes when the concentration increases. Namely, with concentrations ranging from 1.6 to 6.4 μM, the oxidation percentage of FATxPBN equals that of the multiplicative combination of the separated agents, which indicates that FATxPBN exhibits a simple additive effect. At high concentrations (*i.e.* from 12.6 to 38.4 μM), FATxPBN exhibits an infra-additive effect (inhibition between Trolox and PBN), as its oxidation values are higher than the multiplicative combination of FATx and FAPBN. Similar trends (Supp. 5) were observed when using the Chou and Talalay’s method (55) to evaluate synergy, leading to the same conclusions.

Several researchers have reported on synergistic effects created by combining different antioxidant moieties. For instance, it has been reported that the combination of β-carotene and α-tocopherol results in a better protection of rat liver microsomes from oxidative stress (56). Recently, Trolox has been mixed with caffeic acid phenethyl ester (CAPE) to create a synergistic antioxidant effect (57), which was attributed to weak interactions (hydrogen bonds) between Trolox and CAPE. Trolox has also been combined to nitrones to create nitrone derivatives of Trolox (58), which revealed more efficient *in vitro* and *in vivo* activities against lipid peroxidation and ischemia with respect to the parent derivatives. Synergism between antioxidants can proceed through different mechanisms like sacrificial oxidation (59), regeneration or other mechanisms (60). In the case of sacrificial oxidation, one antioxidant protects the other by free radical scavenging, while for regeneration, one antioxidant is oxidized to renew the second one. A famous example of regeneration is the interaction between vitamin E and vitamin C (61). Other researchers attempted to combine, in a single molecule, different types of antioxidants to favor synergistic effects. For instance, lipoic acid has been combined to Trolox (62) as well as a nitric oxide synthase inhibitor (63) with the objective to obtain greater protective effects. Preliminary experiments on biological systems highlighted that these divalent antioxidants have better abilities to prevent neuronal damage caused by glutamate and oxidation of rat microsomal membrane lipids.

Recently, Vavrilokova and co-workers, through enzymatic reactions, combined silybin to ascorbic acid, Trolox alcohol or tyrosol to obtain novel divalent antioxidants (64). Among all the derivatives, the derivative combining silybin to ascorbic acid showed the highest ability to protect rat microsomes from lipid peroxidation. Our group has previously reported the synthesis of PBNLP, a divalent antioxidant bearing a PBN and a lipoic acid moiety (65). No synergistic antioxidant effect was observed as PBNLP and its individual constituents showed a similar ability to protect red blood cells from AAPH-induced hemolysis. However, the combination of the two moieties on the same carrier strongly diminished the antagonist effect observed between the individual derivatives.

In the case of FATxPBN, no intramolecular interactions between both antioxidant (Trolox and PBN) moieties were observed by MD simulations, which preclude regeneration effect within the same molecule. However, regarding to the similar depth of insertion of both Trolox and PBN moieties, intermolecular interactions are likely to occur, which could favor a regeneration process. Also, the ability of FATxPBN to strongly delay lipid oxidation compared to the other antioxidants (Fig. 5) suggests that this derivative combines the retarding effect of PBN (66) with the free radical scavenging capacity of Trolox (67), thus creating a synergistic effect by retardation. At first, the PBN moiety seems to delay the oxidation during 60 min, as percentages of oxidation of liposomes are the same, regardless of the concentration of FATxPBN (Fig. 4A). Then, the dose-dependent antioxidant efficiency observed might be attributed to the Trolox moiety, as FAPBN is only slightly efficient against AAPH-induced oxidation (Fig. 4F).

**Table 3.**
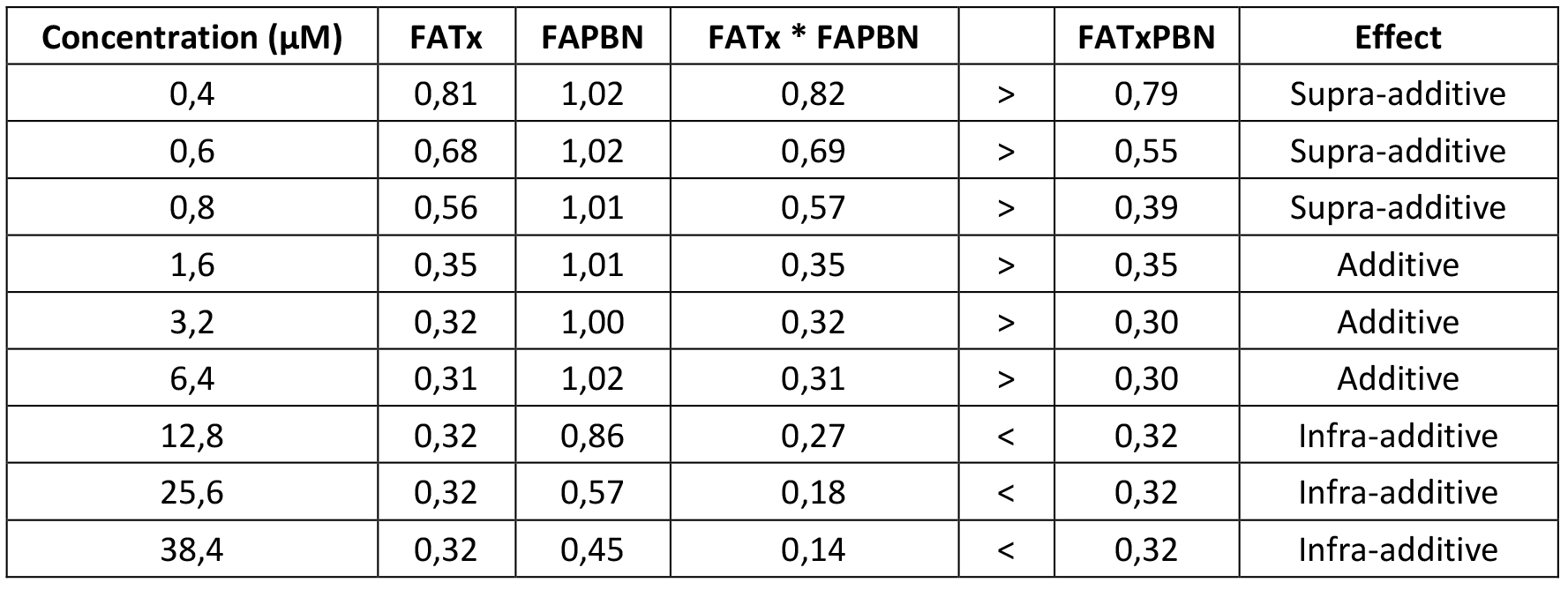
Evaluation of the synergistic effect of FATxPBN. Additive: FATxPBN = FATx * FAPBN. Supra-additive (synergy): FATxPBN < FATx * FAPBN. Infra-additive (antagonist): FATxPBN > FATx * FAPBN.

| Concentration ($\mu M$) | FATx | FAPBN | $FATx \cdot FAPBN$ | | FATxPBN | Effect |
| --- | --- | --- | --- | --- | --- | --- |
| 0,4 | 0,81 | 1,02 | 0,82 | > | 0,79 | Supra-additive |
| 0,6 | 0,68 | 1,02 | 0,69 | > | 0,55 | Supra-additive |
| 0,8 | 0,56 | 1,01 | 0,57 | > | 0,39 | Supra-additive |
| 1,6 | 0,35 | 1,01 | 0,35 | > | 0,35 | Additive |
| 3,2 | 0,32 | 1,00 | 0,32 | > | 0,30 | Additive |
| 6,4 | 0,31 | 1,02 | 0,31 | > | 0,30 | Additive |
| 12,8 | 0,32 | 0,86 | 0,27 | < | 0,32 | Infra-additive |
| 25,6 | 0,32 | 0,57 | 0,18 | < | 0,32 | Infra-additive |
| 38,4 | 0,32 | 0,45 | 0,14 | < | 0,32 | Infra-additive |

## 4. Conclusion

The present work is in the line of the strategy aiming at grafting two antioxidant moieties on the same carrier to enhance the global antioxidant action with a nanoscale fine tuning. The key role of the amphiphilic feature of this series of fluorinated antioxidants bearing Trolox and/or PBN moieties was studied in terms of interaction with membranes. The presence of the perfluorinated chain increases hydrophobicity, allowing these derivatives to form monolayers at the air-water interface. Because of higher hydrophobicity, they are anchored to the DLPC membranes, with a greater antioxidant activity against AAPH-induced lipid oxidation. Interestingly, although PBN is inefficient, its combination with a Trolox moiety on the same carrier creates a synergistic effect that improves the global antioxidant efficiency. Thanks to a dual mode of action, such hybrid antioxidants are very promising molecules for a wide range of applications (cosmetic, pharmaceutical and food) by lowering the concentrations required to efficiently fight against lipid oxidation.

## Contributions

G.D and S.M conceived this research. M.R synthesized the fluorinated derivatives. L.S and A.M.G.G performed the experiments with model membranes. B.C and F.D.M designed and performed MD simulations. B.C, F.D.M and P.T analyzed the theoretical data. L.S analyzed the experimental data and wrote the article. P.T, G.D and S.M formatted the manuscript.

## Acknowledgements

L. Socrier was the recipient of a scholarship from the French “Ministère de l’enseignement supérieur et de la recherche”. Dr Marie Rosselin was the recipient of a fellowship from the “Provence Alpes Côte d’Azur” regional council. The “Centre national de la recherche scientifique (CNRS)”, the “Université de Technologie de Compiègne” and the “Université d’Avignon et des Pays du Vaucluse” are gratefully acknowledged for providing facilities and financial support. P.Trouillas, B. Chantemargue and F. Di Meo thank CALI, “Nouvelle Aquitaine” regional council and the “Institut national de la santé et de la recherche médicale (INSERM)”. P. Trouillas thanks the Czech Science Foundation (P208/12/G016) and National Program of Sustainability from the Ministry of Youth, Education and Sports of the Czech Republic (LO1305).

## References

1. Apel, K., and H. Hirt. 2004. REACTIVE OXYGEN SPECIES: Metabolism, Oxidative Stress, and Signal Transduction. Annual Review of Plant Biology. 55: 373–399.

2. Forman, H.J., and M. Torres. 2002. Reactive oxygen species and cell signaling: respiratory burst in macrophage signaling. American journal of respiratory and critical care medicine. 166: S4–8.

3. Dröge, W. 2002. Free Radicals in the Physiological Control of Cell Function. Physiological Reviews. 82: 47–95.

4. Pincemail, J., M. Meurisse, R. Limet, and J.O. Defraigne. 1999. Espèces oxygénées activées, antioxydants et cancer. Vaisseaux, Coeur, Poumons. 4: 2–5.

5. Sultana, R., M. Perluigi, and D.A. Butterfield. 2013. Lipid peroxidation triggers neurodegeneration: A redox proteomics view into the Alzheimer disease brain. Free Radical Biology and Medicine. 62: 157–169.

6. Kojda, G. 1999. Interactions between NO and reactive oxygen species: pathophysiological importance in atherosclerosis, hypertension, diabetes and heart failure. Cardiovascular Research. 43: 562–571.

7. Circu, M.L., and T.Y. Aw. 2010. Reactive oxygen species, cellular redox systems, and apoptosis. Free Radical Biology and Medicine. 48: 749–762.

8. Waris, G., and H. Ahsan. 2006. Reactive oxygen species role in the development of cancer and various chronic conditions. Journal of Carcinogenesis. 5: 14.

9. Wheatley, R. 2000. Some recent trends in the analytical chemistry of lipid peroxidation. TrAC Trends in Analytical Chemistry. 19: 617–628.

10. Forrest, V.J., Y.-H. Kang, D.E. McClain, D.H. Robinson, and N. Ramakrishnan. 1994. Oxidative stress-induced apoptosis prevented by trolox. Free Radical Biology and Medicine. 16: 675–684.

11. Sagach, V.F., M. Scrosati, J. Fielding, G. Rossoni, C. Galli, and F. Visioli. 2002. The water-soluble vitamin E analogue Trolox protects against ischaemia/reperfusion damage in vitro and ex vivo. A comparison with vitamin E. Pharmacological Research. 45: 435–439.

12. Miller, N.J., C. Rice-Evans, M.J. Davies, V. Gopinathan, and A. Milner. 1993. A Novel Method for Measuring Antioxidant Capacity and its Application to Monitoring the Antioxidant Status in Premature Neonates. Clinical Science. 84: 407–412.

13. van den Berg, R., G.R.M.M. Haenen, H. van den Berg, and A. Bast. 1999. Applicability of an improved Trolox equivalent antioxidant capacity (TEAC) assay for evaluation of antioxidant capacity measurements of mixtures. Food Chemistry. 66: 511–517.

14. Finkelstein, E., G.M. Rosen, and E.J. Rauckman. 1980. Spin trapping of superoxide and hydroxyl radical: Practical aspects. Archives of Biochemistry and Biophysics. 200: 1–16.

15. Floyd, R.A., K. Hensley, M.J. Forster, J.A. Kelleher-Andersson, and P.L. Wood. 2002. Nitrones, their value as therapeutics and probes to understand aging. Mechanisms of Ageing and Development. 123: 1021–1031.

16. Doblas, S., D. Saunders, P. Kshirsagar, Q. Pye, J. Oblander, B. Gordon, S. Kosanke, R.A. Floyd, and R.A. Towner. 2008. Phenyl-tert-butylnitrone induces tumor regression and decreases angiogenesis in a C6 rat glioma model. Free Radical Biology and Medicine. 44: 63–72.

17. Floyd, R.A., H.C.C.F. Neto, G.A. Zimmerman, K. Hensley, and R.A. Towner. 2013. Nitrone-based therapeutics for neurodegenerative diseases: Their use alone or in combination with lanthionines. Free Radical Biology and Medicine. 62: 145–156.

18. Ayuso, M.I., M. Chioua, E. Martínez-Alonso, E. Soriano, J. Montaner, J. Masjuán, D.J. Hadjipavlou-Litina, J. Marco-Contelles, and A. Alcázar. 2015. CholesteroNitrones for Stroke. Journal of Medicinal Chemistry. 58: 6704–6709.

19. Choteau, F., G. Durand, I. Ranchon-Cole, C. Cercy, and B. Pucci. 2010. Cholesterol-based α-phenyl-N-tert-butyl nitrone derivatives as antioxidants against light-induced retinal degeneration. Bioorganic & medicinal chemistry letters. 20: 7405–9.

20. Socrier, L., M. Rosselin, F. Choteau, G. Durand, and S. Morandat. 2017. Cholesterol-nitrone conjugates as protective agents against lipid oxidation: A model membrane study. Biochimica et Biophysica Acta (BBA) - Biomembranes. 1859: 2495–2504.

21. Ortial, S., G. Durand, B. Poeggeler, A. Polidori, M.A. Pappolla, J. Böker, R. Hardeland, and B. Pucci. 2006. Fluorinated amphiphilic amino acid derivatives as antioxidant carriers: A new class of protective agents. Journal of Medicinal Chemistry. 49: 2812–2820.

22. Rosselin, M., G. Meyer, P. Guillet, T. Cheviet, G. Walther, A. Meister, D. Hadjipavlou-Litina, and G. Durand. 2016. Divalent Amino-Acid-Based Amphiphilic Antioxidants: Synthesis, Self-Assembling Properties, and Biological Evaluation. Bioconjugate Chemistry. 27: 772–781.

23. Der Mardirossian, C., M.P. Krafft, T. Gulik-Krzywicki, M. Le Maire, and F. Lederer. 1998. Perfluoroalkylphosphocholines are poor protein-solubilizing surfactants, as tested with neutrophil plasma membranes. Biochimie. 80: 531–541.

24. Fadel, O., K. El Kirat, and S. Morandat. 2011. The natural antioxidant rosmarinic acid spontaneously penetrates membranes to inhibit lipid peroxidation in situ. Biochimica et Biophysica Acta (BBA) - Biomembranes. 1808: 2973–2980.

25. Kondo, M., M. Mehiri, and S.L. Regen. 2008. Viewing membrane-bound molecular umbrellas by parallax analyses. Journal of the American Chemical Society. 130: 13771–13777.

26. Stewart, J.C. 1980. Colorimetric determination of phospholipids with ammonium ferrothiocyanate. Analytical biochemistry. 104: 10–4.

27. Schnitzer, E., I. Pinchuk, and D. Lichtenberg. 2007. Peroxidation of liposomal lipids. European Biophysics Journal. 36: 499–515.

28. Wu, E.L., X. Cheng, S. Jo, H. Rui, K.C. Song, E.M. Dávila-Contreras, Y. Qi, J. Lee, V. Monje-Galvan, R.M. Venable, J.B. Klauda, and W. Im. 2014. CHARMM-GUI membrane builder toward realistic biological membrane simulations. Journal of Computational Chemistry. 35: 1997–2004.

29. Skjevik, Å.A., B.D. Madej, R.C. Walker, and K. Teigen. 2012. LIPID11: A Modular Framework for Lipid Simulations Using Amber. The Journal of Physical Chemistry B. 116: 11124–11136.

30. Wang, J., R.M. Wolf, J.W. Caldwell, P.A. Kollman, and D.A. Case. 2004. Development and testing of a general amber force field. Journal of Computational Chemistry. 25: 1157–1174.

31. Wang, J., and T. Hou. 2011. Application of molecular dynamics simulations in molecular property prediction. 1. Density and heat of vaporization. Journal of Chemical Theory and Computation. 7: 2151–2165.

32. Mobley, D.L., C.I. Bayly, M.D. Cooper, M.R. Shirts, and K.A. Dill. 2009. Small molecule hydration free energies in explicit solvent: An extensive test of fixed-charge atomistic simulations. Journal of Chemical Theory and Computation. 5: 350–358.

33. Kirschner, K.N., A.B. Yongye, S.M. Tschampel, J. González-Outeiriño, C.R. Daniels, B.L. Foley, and R.J. Woods. 2008. GLYCAM06: A generalizable biomolecular force field. carbohydrates. Journal of Computational Chemistry. 29: 622–655.

34. Wang, J., W. Wang, P.A. Kollman, and D.A. Case. 2006. Automatic atom type and bond type perception in molecular mechanical calculations. Journal of Molecular Graphics and Modelling. 25:247–260.

35. Duan, Y., C. Wu, S. Chowdhury, M.C. Lee, G. Xiong, W. Zhang, R. Yang, P. Cieplak, R. Luo, T. Lee, J. Caldwell, J. Wang, and P. Kollman. 2003. A point-charge force field for molecular mechanics simulations of proteins based on condensed-phase quantum mechanical calculations. Journal of Computational Chemistry. 24: 1999–2012.

36. Frish, M.J., G.W. Trucks, H.B. Schlegel, G.E. Scuseria, M.A. Robb, J.R. Cheeseman, G. Scalmani, V. Barone, G.A. Petersson, H. Nakatsuji, X. Li, M. Caricato, A. Marenich, J. Bloino, G. Janesko, R. Gomperts, B. Mennucci, H.P. Hratchian, V. Ortiz, A.F. Izmaylov, J.L. Sonnenberg, D. Williams-Young, F. Ding, F. Lipparini, F. Egidi, J. Goings, B. Peng, A. Petrone, T. Henderson, D. Ranasinghe, V.G. Zakrzewski, J. Gao, N. Rega, G. Zheng, W. Liang, M. Hada, M. Ehara, K. Toyota, R. Fukuda, J. Hasegawa, M. Ishida, T. Nakajima, Y. Honda, O. Kitao, H. Nakai, T. Vreven, K. Throssel, J.A. Montgomery, J.E. Peralta, F. Ogliaro, M. Bearpark, J.J. Heyd, E. Brothers, K.N. Kudin, V.N. Staroverov, T. Keith, R. Kobayashi, J. Normand, K. Raghavachari, A. Rendell, J.C. Burant, S.S. Iyengar, J. Tomasi, M. Cossi, J.M. Millam, M. Klene, C. Adamo, R. Cammi, J.W. Ochterski, R.L. Martin, K. Morokuma, O. Farkas, J.B. Foresman, and D.J. Fox. Gaussian, Inc., Wallingford CT, 2016.

37. Dupradeau, F.-Y., A. Pigache, T. Zaffran, C. Savineau, R. Lelong, N. Grivel, D. Lelong, W. Rosanski, and P. Cieplak. 2010. The R.E.D. Tools: Advances in RESP and ESP charge derivation and force field library building. Physical chemistry chemical physics : PCCP. 12: 7821–7839.

38. Jorgensen, W.L., J. Chandrasekhar, J.D. Madura, R.W. Impey, and M.L. Klein. 1983. Comparison of simple potential functions for simulating liquid water. The Journal of Chemical Physics. 79: 926–935.

39. Darden, T., D. York, and L. Pedersen. 1993. Particle mesh Ewald: An N ⋅ log(N) method for Ewald sums in large systems. The Journal of Chemical Physics. 98: 10089–10092.

40. Case, D.A., D.S. Cerutti, T.E‥ Cheatham, T.A. Darden, R.E. Duke, T.J. Giese, H. Gohlke, A.W. Goetz, D. Greene, and N. Homeyer. AMBER 2017;University of California: San Francisco, CA, USA, 2017‥

41. Salomon-Ferrer, R., D.A. Case, and R.C. Walker. 2013. An overview of the Amber biomolecular simulation package. Wiley Interdisciplinary Reviews: Computational Molecular Science. 3: 198–210.

42. Ryckaert, J.-P., G. Ciccotti, and H.J.C. Berendsen. 1977. Numerical integration of the cartesian equations of motion of a system with constraints: molecular dynamics of n-alkanes. Journal of Computational Physics. 23: 327–341.

43. Loncharich, R.J., B.R. Brooks, and R.W. Pastor. 1992. Langevin dynamics of peptides: The frictional dependence of isomerization rates of N-acetylalanyl-N′-methylamide. Biopolymers. 32: 523–535.

44. Berendsen, H.J.C., J.P.M. Postma, W.F. Van Gunsteren, A. Dinola, and J.R. Haak. 1984. Molecular dynamics with coupling to an external bath. The Journal of Chemical Physics. 81: 3684–3690.

45. Roe, D.R., and T.E. Cheatham. 2013. PTRAJ and CPPTRAJ: Software for Processing and Analysis of Molecular Dynamics Trajectory Data. Journal of Chemical Theory and Computation. 9: 3084–3095.

46. Calvez, P., S. Bussières, Éric Demers, and C. Salesse. 2009. Parameters modulating the maximum insertion pressure of proteins and peptides in lipid monolayers. Biochimie. 91: 718–733.

47. Marsh, D. 1996. Lateral pressure in membranes. Biochimica et Biophysica Acta (BBA) - Reviews on Biomembranes. 1286: 183–223.

48. Frotscher, E., B. Danielczak, C. Vargas, A. Meister, G. Durand, and S. Keller. 2015. A Fluorinated Detergent for Membrane-Protein Applications. Angewandte Chemie International Edition. 54: 5069–5073.

49. Marquardt, D., J.A. Williams, N. Kučerka, J. Atkinson, S.R. Wassall, J. Katsaras, and T.A. Harroun. 2013. Tocopherol activity correlates with its location in a membrane: A new perspective on the antioxidant vitamin e. Journal of the American Chemical Society. 135: 7523–7533.

50. Garrec, J., A. Monari, X. Assfeld, L.M. Mir, and M. Tarek. 2014. Lipid peroxidation in membranes: The peroxyl radical does not “float.” Journal of Physical Chemistry Letters. 5: 1653–1658.

51. Košinová, P., K. Berka, M. Wykes, M. Otyepka, and P. Trouillas. 2012. Positioning of Antioxidant Quercetin and Its Metabolites in Lipid Bilayer Membranes: Implication for Their Lipid-Peroxidation Inhibition. The Journal of Physical Chemistry B. 116: 1309–1318.

52. Azouzi, S., H. Santuz, S. Morandat, C. Pereira, F. Côté, O. Hermine, K. El Kirat, Y. Colin, C. Le Van Kim, C. Etchebest, and P. Amireault. 2017. Antioxidant and Membrane Binding Properties of Serotonin Protect Lipids from Oxidation. Biophysical Journal. 112: 1863–1873.

53. Berenbaum, M.C. 1981. Criteria for analyzing interactions between biologically active agents. Advances in Cancer Research. 35: 269–335.

54. Hennequin, C., N. Giocanti, and V. Favaudon. 1996. Interaction of ionizing radiation with paclitaxel (taxol) and docetaxel (taxotere) in HeLa and SQ20B cells. Cancer Research. 56: 1842–1850.

55. Chou, T.C., and P. Talalay. 1984. Quantitative analysis of dose-effect relationships: the combined effects of multiple drugs or enzyme inhibitors. Advances in Enzyme Regulation. 22: 27–55.

56. Palozza, P., and N.I. Krinsky. 1992. Beta-Carotene and alpha-tocopherol are synergistic antioxidants. Archives of Biochemistry and Biophysics. 297: 184–187.

57. Bai, H., R. Liu, H.L. Chen, W. Zhang, X. Wang, X. Di Zhang, W.L. Li, and C.X. Hai. 2014. Enhanced antioxidant effect of caffeic acid phenethyl ester and Trolox in combination against radiation induced-oxidative stress. Chemico-Biological Interactions. 207: 7–15.

58. Balogh, G.T., K. Vukics, Á. Könczöl, Á. Kis-Varga, A. Gere, and J. Fischer. 2005. Nitrone derivatives of trolox as neuroprotective agents. Bioorganic & Medicinal Chemistry Letters. 15: 3012–3015.

59. Neunert, G., P. Górnaś, K. Dwiecki, A. Siger, and K. Polewski. 2015. Synergistic and antagonistic effects between alpha-tocopherol and phenolic acids in liposome system: spectroscopic study. European Food Research and Technology. 241: 749–757.

60. Becker, E.M., L.R. Nissen, and L.H. Skibsted. 2004. Antioxidant evaluation protocols: Food quality or health effects. European Food Research and Technology. 219: 561–571.

61. Kamal-Eldin, A., and L.-Å. Appelqvist. 1996. The chemistry and antioxidant properties of tocopherols and tocotrienols. Lipids. 31: 671–701.

62. Koufaki, M., T. Calogeropoulou, A. Detsi, A. Roditis, A.P. Kourounakis, P. Papazafiri, K. Tsiakitzis, C. Gaitanaki, I. Beis, and P.N. Kourounakis. 2001. Novel potent inhibitors of lipid peroxidation with protective effects against reperfusion arrhythmias. Journal of Medicinal Chemistry. 44: 4300–4303.

63. Harnett, J.J., M. Auguet, I. Viossat, C. Dolo, D. Bigg, and P.E. Chabrier. 2002. Novel lipoic acid analogues that inhibit nitric oxide synthase. Bioorganic & medicinal chemistry letters. 12: 1439–1442.

64. Vavříková, E., V. Křen, L. Jezova-Kalachova, M. Biler, B. Chantemargue, M. Pyszková, S. Riva, M. Kuzma, K. Valentová, J. Ulrichová, J. Vrba, P. Trouillas, and J. Vacek. 2017. Novel flavonolignan hybrid antioxidants: From enzymatic preparation to molecular rationalization. European Journal of Medicinal Chemistry. 127: 263–274.

65. Durand, G., A. Polidori, J.P. Salles, M. Prost, P. Durand, and B. Pucci. 2003. Synthesis and antioxidant efficiency of a new amphiphilic spin-trap derived from PBN and lipoic acid. Bioorganic and Medicinal Chemistry Letters. 13: 2673–2676.

66. Barclay, L.R.C., and M.R. Vinqvist. 2000. Do spin traps also act as classical chain-breaking antioxidants? A quantitative kinetic study of phenyl tert-butylnitrone (PBN) in solution and in liposomes. Free Radical Biology and Medicine. 28: 1079–1090.

67. Franzoni, F., R. Colognato, F. Galetta, I. Laurenza, M. Barsotti, R. Di Stefano, R. Bocchetti, F. Regoli, A. Carpi, A. Balbarini, L. Migliore, and G. Santoro. 2006. An in vitro study on the free radical scavenging capacity of ergothioneine: comparison with reduced glutathione, uric acid and trolox. Biomedicine & Pharmacotherapy. 60: 453–457.

